# Powassan Viruses Spread Cell to Cell During Direct Isolation from *Ixodes* Ticks and Persistently Infect Human Brain Endothelial Cells and Pericytes

**DOI:** 10.1101/2021.09.30.462684

**Authors:** Jonas N. Conde, Santiago Sanchez-Vicente, Nicholas Saladino, Elena E. Gorbunova, William R. Schutt, Megan C. Mladinich, Grace Himmler, Jorge Benach, Hwan Keun Kim, Erich R Mackow

## Abstract

Powassan viruses (POWVs) are neurovirulent tick-borne flaviviruses emerging in the Northeastern U.S., with a 2% prevalence in Long Island (LI) deer ticks (*Ixodes scapularis*). POWVs are transmitted in as little as 15 minutes of a tick bite, and enter the CNS to cause encephalitis (10% fatal) and long-term neuronal damage. POWV-LI9 and POWV-LI41 present in LI *Ixodes* ticks were isolated by directly inoculating VeroE6 cells with tick homogenates and detecting POWV infected cells by immunoperoxidase staining. Inoculated POWV-LI9 and LI41 were exclusively present in infected cell foci, indicative of spread cell to cell, despite growth in liquid culture without an overlay. Cloning and sequencing establish POWV-LI9 as a phylogenetically distinct lineage II POWV strain circulating in LI deer ticks. Primary human brain microvascular endothelial cells (hBMECs) and pericytes form a neurovascular complex that restricts entry into the CNS. We found that POWV-LI9, -LI41 and Lineage I POWV-LB, productively infect hBMECs and pericytes and that POWVs were basolaterally transmitted from hBMECs to lower chamber pericytes without permeabilizing polarized hBMECs. Synchronous POWV-LI9 infection of hBMECs and pericytes induced proinflammatory chemokines, interferon-β (IFNβ) and IFN-stimulated genes, with delayed IFNβ secretion by infected pericytes. IFN inhibited POWV infection, but despite IFN secretion a subset of POWV infected hBMECs and pericytes remained persistently infected. These findings suggest a potential mechanism for POWVs (LI9/LI41 and LB) to infect hBMECs, spread basolaterally to pericytes and enter the CNS. hBMEC and pericyte responses to POWV infection suggest a role for immunopathology in POWV neurovirulence and potential therapeutic targets for preventing POWV spread to neuronal compartments.

**Importance:** We isolated POWVs from LI deer ticks (*I. scapularis*) directly in VeroE6 cells and sequencing revealed POWV-LI9 as a distinct lineage II POWV strain. Remarkably, inoculating VeroE6 cells with POWV containing tick homogenates resulted in infected cell foci in liquid culture, consistent with cell to cell spread. POWV-LI9, -LI41, and Lineage I POWV-LB strains infected hBMECs and pericytes that comprise neurovascular complexes. POWVs were nonlytically transmitted basolaterally from infected hBMECs to lower chamber pericytes, suggesting a mechanism for POWV transmission across BBB. POWV-LI9 elicited inflammatory responses from infected hBMEC and pericytes that may contribute to immune cell recruitment and neuropathogenesis. This study reveals a potential mechanism for POWVs to enter the CNS by infecting hBMECs and spreading basolaterally to abluminal pericytes. Our findings reveal that POWV-LI9 persists in cells that form a neurovascular complex spanning the BBB, and suggest potential therapeutic targets for preventing POWV spread to neuronal compartments.

## Introduction

Powassan virus (POWV) is a neurovirulent tick-borne flavivirus (FV) that was first isolated in 1958 from a fatal encephalitic case of human disease in Powassan, Ontario, Canada^(1)^. POWV has recently emerged in ticks in the Northeastern U.S., with host reservoirs in white-footed mice, deer, skunks, woodchucks, and squirrels^(2–5)^. In humans, POWV enters the central nervous system (CNS), lytically infects neurons, and causes severe encephalitis with a ~10% mortality rate and long-term neurologic sequelae occurring in 50% of infected patients^(6–9)^. POWV awareness and diagnostic testing of patients has revealed an increasing number of human POWV encephalitis cases that suggest a high incidence of previously undiagnosed POWV disease, especially in areas where Lyme disease is endemic. Approximately 2% of deer ticks *(I. scapularis)*, in the highly populated Long Island (LI), N.Y. area are POWV positive^(6)^. Currently there are no approved vaccines or therapeutics for POWV.

The presence of POWV in tick saliva accounts for POWV transmission to hosts or humans in as little as 15 minutes after a tick bite^(8–10)^. POWV transmission is followed by a 1-5 week incubation period, a 1 week of acute febrile illness and weeks to months of encephalitic manifestations of fever, headache and confusion that worsen into meningoencephalitis with seizures, impaired consciousness, coma and respiratory failure^(9)^. In autopsies of POWV patients, neurons in the central nervous system (CNS) were infected with POWV antigen and RNA within lesions in the cerebellum, brainstem, basal ganglia, and thalamus^(11)^. The emergence of neurovirulent POWV in highly populated areas, rapid tick to human transmission and the severity of POWV disease provide urgency for understanding how POWVs spread to neuronal compartments, and defining potential therapeutic targets.

POWV is the only North American tick-borne flavivirus and POWVs are distantly related to Tick Borne Encephalitis virus (TBEV), the leading cause of arthropod borne encephalitis in Europe and Asia^(12)^. There are two genetic lineages of POWV, lineage I includes the LB strain derived from the 1958 human encephalitis case, and lineage II strains sequenced or isolated from North American deer ticks. POWV I and II strains share 94% amino acid identity and represent POWV genotypes carried by discrete ticks and mammalian hosts^(13)^. Prototype Lineage I and II POWVs were first isolated following intracranial inoculation of mice and subsequent passage in Vero cells^(1, 8, 14, 15)^. However, there are no studies of POWV infection of cells that comprise the BBB and normally prevent viral access to neuronal compartments.

How POWV spreads from a tick bite site to the CNS, persists to cause a late acute febrile illness, or enters protected neuronal compartments remains to be determined. Reservoir cells where POWV persists following infection and mechanisms by which POWV crosses the BBB to enter the CNS remain enigmatic. There are no reports of POWV infection of hBMECs or pericytes, or cellular responses to POWV or TBEV infections. TBEV was found to infect only 2-5% of human brain microvascular ECs (hBMECs), was primarily apically released, and failed to lyse ECs, alter EC barrier functions or spread within EC monolayers^(16)^. In mice, TBEV reportedly enters the CNS without permeabilizing the BBB, and at late stages of infection induces proinflammatory cytokine and chemokine responses when high viral loads are already present in the brain^(17)^.

hBMECs have intimate basolateral contacts with pericytes, forming a neurovascular unit that plays a critical role in regulating BBB permeability and immune cell recruitment. POWV infection of hBMECs and brain pericytes have not been investigated. In this study, we isolated POWVs from Long Island deer ticks and evaluated their ability to replicate, persistently infect, and spread in VeroE6 cells, primary human BMECs and brain pericytes. We collected 438 deer ticks from Long Island, and from 44 pools of 10 ticks, we found 3 POWV positive pools by qRT-PCR. Tick homogenates inoculated into 44 wells of VeroE6 cells revealed 2 wells (9 and 41) with POWV antigen positive cells 7 dpi, both coinciding with PCR positivity. We observed that direct inoculation of VeroE6 cells with LI tick homogenates resulted in large POWV infected cell foci consistent with cell to cell spread despite the absence of restrictive overlays. POWV-LI9 and POWV-LI41 were propagated from supernatants and POWV-LI9 was cloned and sequenced, identifying it as a distinct Lineage II POWV strain.

We found that POWV-LI9 and POWV-LI41 strains productively and persistently infected VeroE6 cells, hBMECs and human pericytes nonlytically, reaching respective POWV-LI9 titers of 8 x 10^5^, 1 x 10^4^, and 5 x10^4^. POWV-LI9 failed to permeabilize polarized hBMECs or VeroE6 cells, but was preferentially released basolaterally from cells (3-12-fold) to lower chamber pericytes in a BBB-like Transwell model. Prototypic POWV-LB acquired from the ATCC has been passaged extensively in Vero cells since 1958^(1)^, and similar analysis of POWV-LB infection of VeroE6 cells resulted in a few small clusters of infected cells but primarily spread uniformly within the monolayer. However, like POWV-LI9, we found that POWV-LB also productively infected hBMECs and pericytes and POWV-LB was basolaterally released from polarized hBMECs (3-fold over apical titers). These findings suggest a potential mechanism for Lineage I and II POWVs to infect hBMECs and pericytes and cross the BBB.

Lectin pretreating hBMECs permitted POWV-LI9 to synchronously infect hBMECs (>90%) and we used this approach to evaluate hBMEC and pericyte responses to POWV infection. POWV induced IFNβ, CCL5 (>3000-fold) an array of inflammatory chemokines, and IFN-stimulated genes (ISGs) 1-3 dpi, with conserved and novel responses between POWV infected hBMECs and pericytes. IFNβ induction and secretion were restricted in POWV infected pericytes, but not hBMECs, 1 dpi, but IFNβ was highly secreted by both hBMECs and pericytes 2 dpi. Pretreating VeroE6 cells, hBMECs or pericytes with Type I IFN prevented POWV infection. Despite IFNβ induction and sensitivity, a subset of hBMECs and pericytes were persistently infected with POWV-LI9, 12-30 dpi and through cell passage. These findings differ from ZIKV which spreads and replicates continuously in hBMECs in the absence of secreted IFNβ^(18)^. Our results reveal novel POWV-LI9 regulation of hBMEC and pericyte responses and suggest that hBMEC-pericyte neurovascular complexes may serve as POWV reservoirs and conduits for POWV to spread basolaterally across the BBB and into neuronal compartments.

## Results

### POWV-LI9 isolation from *Ixodes scapularis* ticks

In November 2020 we collected 438 *I. scapularis* adult ticks, in Suffolk County, Long Island, New York. Homogenates of 44 pools of 10 ticks were prepared in PBS by dounce homogenization, and following centrifugation, supernatants were inoculated into VeroE6 cells and hBMECs in 24-well plates and screened for POWV RNA by qRT-PCR. Three tick pool inocula were PCR positive for POWV RNA. One week after inoculation, infected cell supernatants were harvested and cell monolayers were immunoperoxidase stained using anti-POWV HMAF antibody (ATCC). Coincident with RNA positive tick pools, VeroE6 cell wells 9 and 41 had POWV antigen positive infected cells 7 dpi (Figure 1A).

**Figure 1.**
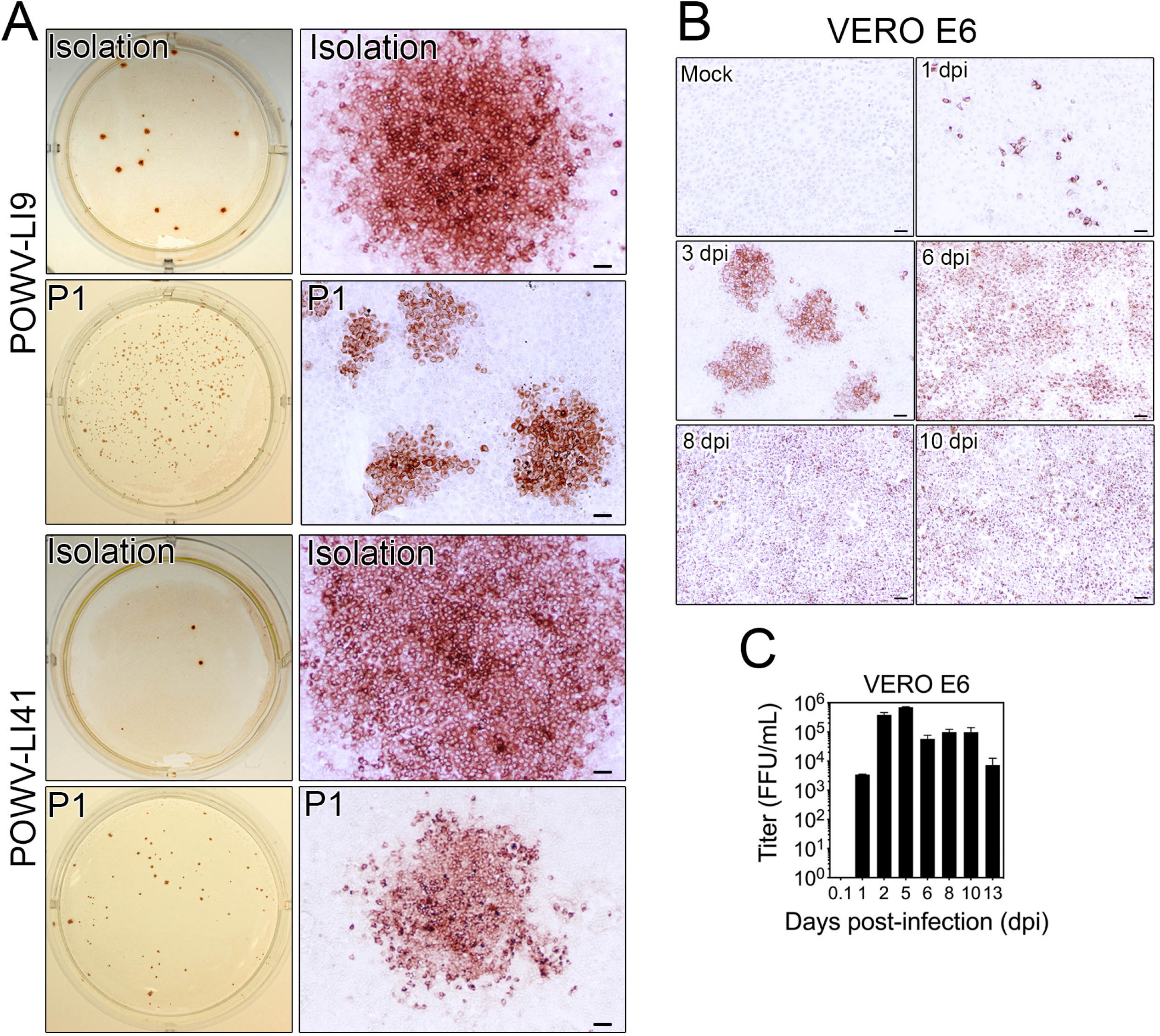
POWV-LI9 isolation from *Ixodes scapularis* ticks. (A) *I. scapularis* ticks were collected in Long Island, N.Y. Tick homogenates from groups of 10 ticks were added to wells of 24-well plates containing VeroE6 cells. After 7 days, cells were fixed and immunoperoxidase stained using a specific anti-POWV HMAF (1:10,000). Wells #9 and # 41 were positive for POWV antigen and formed distinct infected cell foci. First passage (P1) virus was inoculated into a well of 24-well plate, and cells were fixed and immunoperoxidase stained using a specific anti-POWV HMAF (1:10,000) after 7 days (left panels). Right panels show the magnification of the cell foci from initial viral isolation and P1 infection of VeroE6 cells. Bars represent 50 μm. (B) VeroE6 were infected with POWV-LI9 (P3) at an MOI of 5 or mock-infected, cells were fixed at 1, 3, 5, 8, and 10 dpi, and immunoperoxidase stained using anti-POWV HMAF (1:10,000). Bars represent 50 μm. (C) Viral titration from POWV-infected VeroE6 supernatants was determined by FFU assay at 1 dpi in Vero E6 cells.

Despite growth in liquid culture, POWV antigen positive VeroE6 cells were present in large infected cell foci, consistent with cell to cell spread, without apparent cell lysis (Figure 1A). Infection of VeroE6 cells with passage 1 POWV-LI9 or POWV-LI41 stocks resulted in smaller more dispersed infected cell foci (Figure 1A). A time course of POWV-LI9 spread in VeroE6 cells from 1-10 dpi shows that virus spread from initial infected cell foci 3-5 dpi to uniformly infect monolayers 6-10 dpi (Figure 1B). Similar initial foci and later spread was observed when POWV-LI41 infected VeroE6 cells (Supp. Fig. 1). There was no evidence of cytopathic effects (CPE) in POWV-LI9 infected VeroE6 cells with ~100% of VeroE6 cells still infected 10-13 dpi (Figure 1B). Supernatant titers of POWV-LI9 in VeroE6 cells reached 8 x 10^5^ FFU/ml (Figure 1C) and VeroE6 cells remained highly infected following cellular passage (>10^5^ FFU/mL). Infection of VeroE6 cells with extensively passaged Lineage I POWV-LB (ATCC-1958) resulted in some small clusters of 3-12 infected VeroE6 cells 1 dpi (Supp. Fig. 2A), and the uniform infection of VeroE6 cell monolayers with titers of 1 x 10^6^/ml (Supp. Fig. 2A-B). As POWV spread in tissue culture cells following direct tick inoculation has not been previously examined, it remains to be seen if initial cell to cell spread is unique to POWV-LI9/LI41 or a fundamental attribute of direct tick inoculation of POWVs in mammalian cells. Overall these findings demonstrate the direct isolation and productive replication of POWV-LI9 and POWV-LI41 from LI ticks in VeroE6 cells.

### POWV-LI9 is a Discrete Lineage II Strain of POWV Present in LI Deer Ticks

The POWV-LI9 isolate was passaged in VeroE6 cells for 7-10 days with supernatants collected for viral stocks and we cloned and sequenced the POWV-LI9 genome from RNA isolated from passage 3 POWV-LI9 infected VeroE6 cell lysates. Random primed cDNAs were amplified into ~2 kb fragments, using conserved POWV primers, and cloned into pMiniT 2.0. Clones encompassing the complete POWV genome were sequenced with 5’ RACE and a 3’ end oligo used to amplify non-coding POWV-LI9 genome termini (GeneBank accession MZ576219)^(19)^.

Sequences of POWV-LI9 were phylogenetically compared to 15 full-length POWV lineage II sequences, 10 POWV lineage I sequences and TBEV^(20, 21)^. Phylogenetic tree analysis revealed that the POWV-LI9 isolate is a lineage II POWV strain that forms a distinct clade by comparison with other Northeastern, New York and Wisconsin POWV lineage II sequences (Figure 2A)^(20–22)^. Sequence alignment of POWV strains with POWV-LI9 revealed an average 99% nucleotide and amino acid identity with POWV genomes collected in the Northeastern region (Figure 2B, group A), 93.5% nucleotide and 97% amino acid identity with POWV collected in Wisconsin (Figure 2B, group B), and 85% nucleotide and 94.7% amino acid identity with POWV Lineage I (Figure 2B, group C). Novel POWV–LI9 residues include Envelope (Env) protein residues S119 and K151, and NS2B (F125), NS3 (D402) and NS5 (T282) residues. The POWV-LI41 isolate is derived from discrete tick homogenates but has yet to be sequenced to determine if it differs from POWV-LI9.

**Figure 2.**
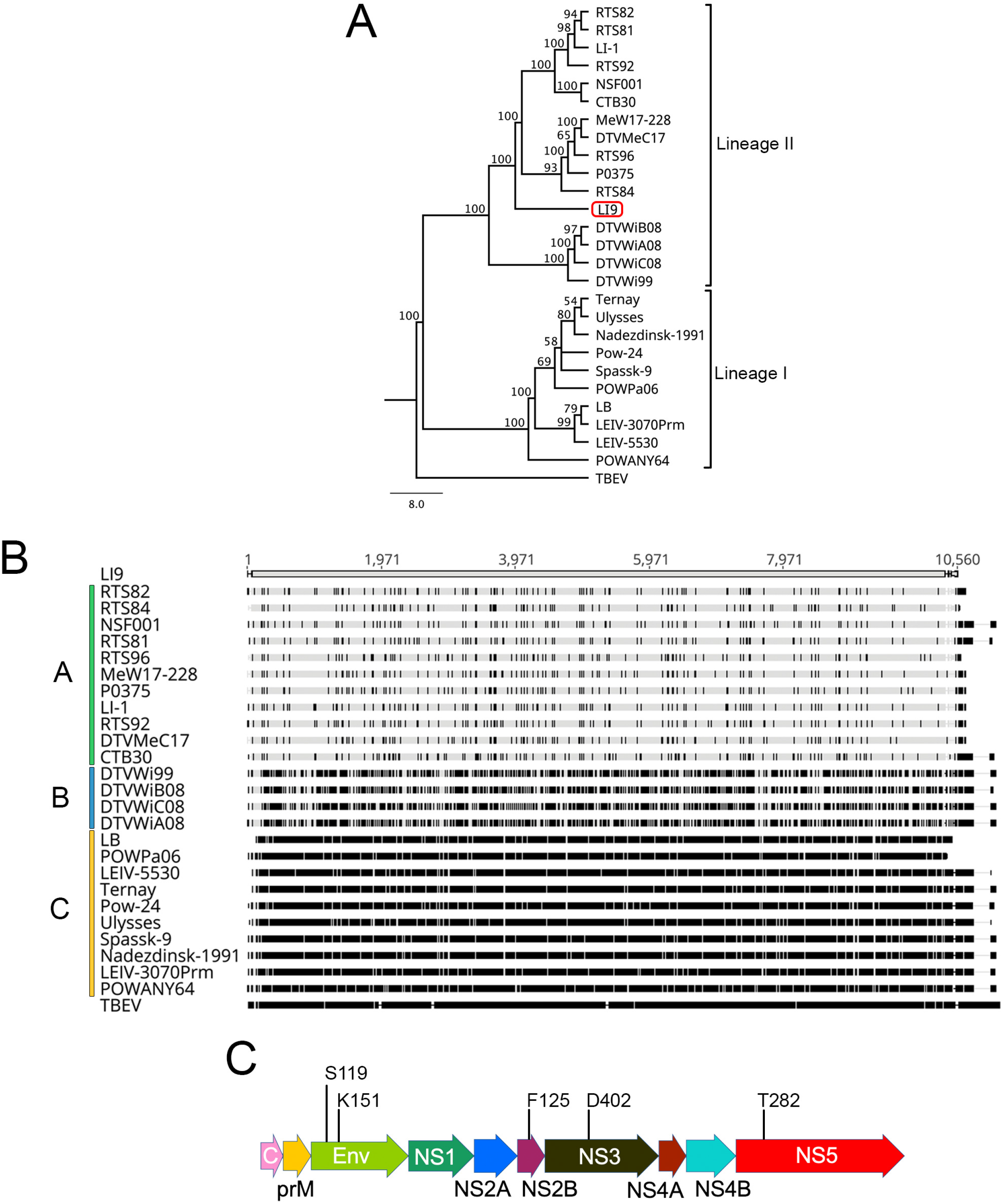
Genetic Analysis of POWV-LI9 Isolates. (A) Maximum-likelihood tree analysis between POWV-LI9 and POWV-related strains using complete genome sequences^(84)^. Branch numbers are bootstrap confidence estimates based on 100 replicates. Tick-borne encephalitis virus (TBEV) is shown as an outgroup. (B) Multiple-sequence analysis comparison using Clustal Omega^(87)^ between POWV-LI9 and other POWV sequences deposited in the GenBank. POWV genomes were divided into three groups based on the genetic identity compared with POWV-LI9: sequences from the viruses collected in the Northeastern U.S. (group A), in Wisconsin (group B), and sequences from lineage I POWV (group C). Black bars within the sequences represent nucleotide differences compared to POWV-LI9. (C) Among 5 residue differences with Northeastern POWVs there are 2 not conserved residues in the Envelope (Env), including Domain II residues S119 and K151 in POWV-LI9 and additional unique residues in the NS2B (F125), NS3 (D402) and NS5 (T282) proteins of POWV-LI9.

In initial experiments, POWV-LI9 (10^3^ FFUs) was inoculated into the footpad of C57BL/6 mice and sera collected 12 dpi were found to elicit antibody responses that detect envelope proteins from POWV-LI9 and POWV-LB infected VeroE6 cells by Western blot (Supp. Fig. 3A); and neutralize POWV-LI9 and POWV-LB viruses (Supp. Fig. 3B). Collectively, these findings establish that POWV-LI9 is a lineage II POWV strain newly discovered in *Ixodes s*. ticks on Long Island, that infects mice and elicits neutralizing antibody responses to Lineage I and II POWVs.

### POWVs Productively Infect Primary Human BMECs and Pericytes

Mechanisms by which POWVs enter neuronal compartments in order to lytically infect neurons have not been investigated. Here we assessed the ability of POWV-LI9 to infect, replicate and persist in primary human BMECs and pericytes that form a neurovascular complex within the BBB^(23–26)^. We found that POWV-LI9 productively infects hBMECs 1-8 dpi, with low viral titers and a subset of hBMECs remaining persistently infected 6-8 dpi (Figure 3A,B). Infecting hBMECs with POWV-LI9 at an MOI of 5 only marginally increased the number of infected hBMECs 1 dpi (10-15% max), suggesting that entry was restricted to a subset of hBMECs.

**Figure 3.**
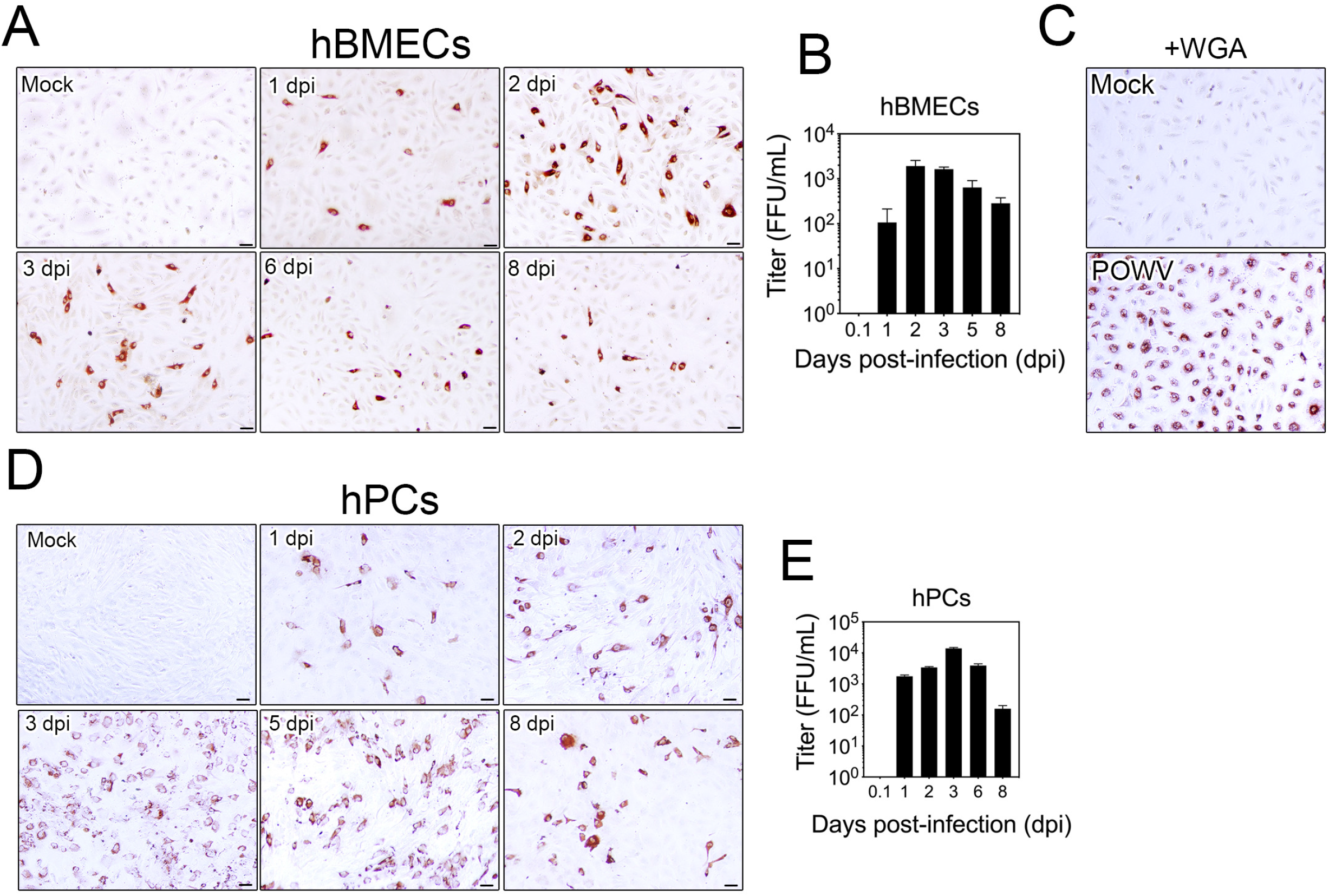
POWV-LI9 Productively and Persistently Infect human BMECs and Pericytes. (A) hBMECs were infected with POWV-LI9 (P3) at an MOI of 5 or mock-infected, cells were fixed at 1, 2, 3, 6, and 8 dpi, and immunoperoxidase stained using anti-POWV HMAF (1:10,000). Bars represent 50 μm. (B) Viral titration from POWV-infected hBMECs supernatants was determined by FFU assay at 1 dpi in Vero E6 cells. (C) POWV-LI9 was pretreated with WGA (1 μg/mL) for 5 min. Control mock WGA treated media (1 μg/mL), or WGA treated POWV was adsorbed to hBMECs for 1 hour, and subsequently cells were PBS washed and resupplemented with EBM2-2% FBS media. Cells were fixed and immunoperoxidase stained using anti-POWV HMAF (1:10,000) at 1 dpi. (D) Human brain pericytes (hPCs) were infected with POWV-LI9 (P3) at an MOI of 5 or mock-infected, cells were fixed at 1, 2, 3, 6, and 8 dpi, and immunoperoxidase stained using a specific anti-POWV HMAF (1:10,000). Bars represent 50 μm. (E) Viral titers from POWV-infected pericyte supernatants were determined by FFU assay at 1 dpi in Vero E6 cells.

To determine if attachment limited hBMEC infection, we treated POWV-LI9 with wheat germ agglutinin (WGA, 1 μg/ml, 5 min) prior to infecting hBMECs, a method previously used to direct efficient HIV infection of ECs^(27–29)^. Pretreating POWV with WGA prior to adsorbing POWV-LI9 to cells, resulted in a synchronous ~95% infection of hBMECs (1 dpi) (Figure 3C). The ability of WGA treated POWV to efficiently direct the infection of hBMECs suggests that POWV attachment and entry, rather than post-attachment replication, is restricted to a subset of hBMECs and permits analysis of hBMEC responses to a synchronous POWV infection.

In BBB capillaries, human brain pericytes are attached to the basolateral side of hBMECs and form abluminal contacts with astrocytes and neurons ^(23–25, 30, 31)^. Pericyte responses and interactions with hBMECs control the integrity and immune cell recruiting functions of the BBB^(31)^. We found that POWV-LI9 infected primary human brain pericytes spreading from 1 to 3 dpi to infect ~80% of pericytes (Figure 3D) and produce titers of 5 x 10^4^ /ml (Figure 3E) without apparent cytopathology. From 3-8 dpi POWV titers were reduced in pericyte supernatants, but a high percentage of pericytes remained POWV infected 8-30 dpi and after pericyte passage. Unlike POWV-LI9 infection of VeroE6 cells, we found that POWV-LI9 infection of pericytes failed to form infected cell foci and instead infected single cells that spread ubiquitously. Similar to POWV-LI9 findings, we found that POWV-LI41 and POWV-LB also productively infect hBMECs and pericytes (Supp. Fig. 4). Collectively, these findings demonstrate that Lineage I and II POWVs infect hBMECs and pericytes which form neurovascular complexes within the BBB, and thereby suggest potential mechanisms for POWVs to enter neuronal compartments.

### Preferential Basolateral Release of POWVs from hBMECs Directs Pericyte Infection

hBMECs form polarized monolayers with apical and basolateral surfaces that mimic luminal and abluminal capillary surfaces^(24)^. The ability of POWV-LI9 to infect hBMECs prompted us to determine whether POWV spreads basolaterally across polarized hBMECs. Neither POWV infected hBMECs, pericytes, or VeroE6 cells showed signs of cytopathic effects and 3 dpi POWV-LI9 infected cells were nearly all viable by Calcein-AM staining (green-live) with little propidium iodide uptake (red-dead) (Figure 4A). Consistent with this, we found no apparent disruption of tight junction protein ZO-1 in cell-to-cell contacts of POWV-LI9 infected hBMECs (Figure 4B).

**Figure 4.**
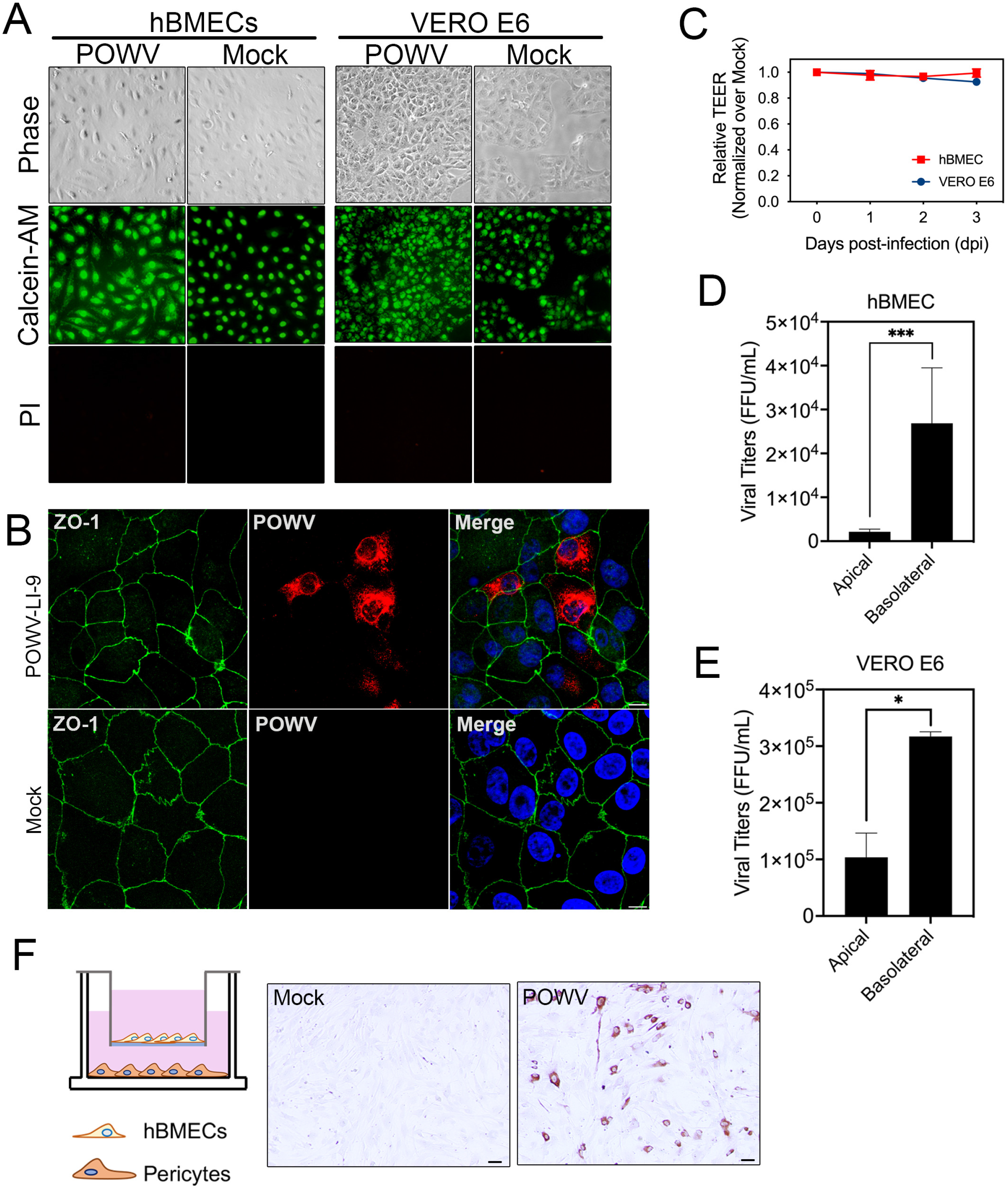
POWV-LI9 is Basolaterally Released from Cells Without Barrier Disruption. (A) Polarized hBMECs and VeroE6 cells grown for 5 days in Transwell inserts, were apically infected with POWV-LI9 (MOI 5). Cells were visualized by phase contrast microscopy for CPE and costained with Calcein-AM (green-live cells) and propidium iodide (red-dead cells) to visualize cell viability 3 dpi. (B) Confocal immunofluorescence. VeroE6 grown on microslides were infected with POWV-LI9 (MOI, 5). Cells were fixed 2 dpi, immunostained with anti-POWV HMAF and anti-ZO-1 antibodies, DAPI counterstained, and visualized by confocal microscopy. Experiments were done in triplicate, repeated at least three times, and representative data are presented. Bars represent 10 μm. (C) Polarized hBMECs and VeroE6 cells grown for 5 days in Transwell inserts, were apically infected with POWV-LI9 (MOI, 5) in triplicate, and TEER was measured 1 to 3 dpi. (D) hBMECs and (E) VeroE6, grown for 5 days in Transwell inserts, were apically infected with POWV-LI9 (MOI, 5) in triplicate. Cells were washed and viral titers from both apical or basolateral compartments were titered 3 dpi. (F) Schematic representation of hBMECs and human pericyte co-culture model (left); hBMECs were seeded on the apical side of Transwell inserts, grown for 7 days confluent by TEER resistance. hBMECs were mock or POWV infected from the apical side for 1 h, cells were washed, and 1 dpi Transwell inserts were transferred to 24-well plates containing monolayers of primary human pericytes. Pericytes were immunoperoxidase stained for POWV antigen 3 dpi using anti-POWV HMAF (1:10,000). Bars represent 50 μm.

To determine whether POWVs permeabilize hBMECs or VeroE6 cells we analyzed polarized cells grown on Transwell inserts for changes in Trans endothelial electrical resistance (TEER). We found no significant change in TEER of POWV infected VeroE6 or hBMECs 1-3 dpi versus uninfected cells, indicating that POWV infection did not permeabilize infected hBMEC monolayers (Figure 4C). Infection of hBMECs and VeroE6 cells on polarized Transwell inserts permitted analysis of apical and basolateral titers following POWV infection. We found that POWV-LI9 titers in basolateral supernatants were enhanced >3-fold in VeroE6 and >12-fold in hBMECs over apically released virus (Figure 4D, E). Analysis of POWV-LB found that POWV titers were enhanced 3 fold in basolateral versus apical supernatants (Supp. Fig. 4D).

POWV basolateral release from hBMECs suggested the potential for POWV to directly infect pericytes. We POWV-LI9 infected polarized hBMECs on Transwell inserts, transferred inserts to wells containing lower chamber pericytes (1dpi) and found that pericytes were infected by basolaterally released POWV (Figure 4F). Preferential basolateral release of POWV from hBMECs suggests a potential mechanism for POWVs to spread across the BBB to abluminal pericytes that reside within neuronal compartments.

### Interferon Pretreatment blocks POWV Infection and Spread

To determine if POWV infections are IFN sensitive, we pretreated VeroE6 cells, hBMECs and pericytes with Type I IFN (IFNα) 2-18 hours prior to POWV-LI9 infection and quantitated infected cells 1 dpi. We found that pretreating cells with IFNα nearly completely abolished POWV infection (Figure 5A), indicating that preexisting or induced Type I IFN prevents POWV infection and spread in hBMECs and pericytes. We analyzed POWV infected hBMEC and pericyte supernatants for IFNβ by ELISA and found that POWV infected hBMECs highly secreted IFNβ 1 dpi, while IFNβ secretion by POWV infected pericytes was not apparent until 2 dpi (Figure 5B,C). Secreted IFNβ responses are consistent with the early restriction of POWV-LI9 spread in hBMECs, and continued POWV spread in pericytes 1-3 dpi (Figure 5D). In both cell types IFNβ secretion is dramatically reduced 3 dpi suggesting a potential mechanism for POWVs to inhibit late IFN expression and persist in infected hBMECs and pericytes.

**Figure 5.**
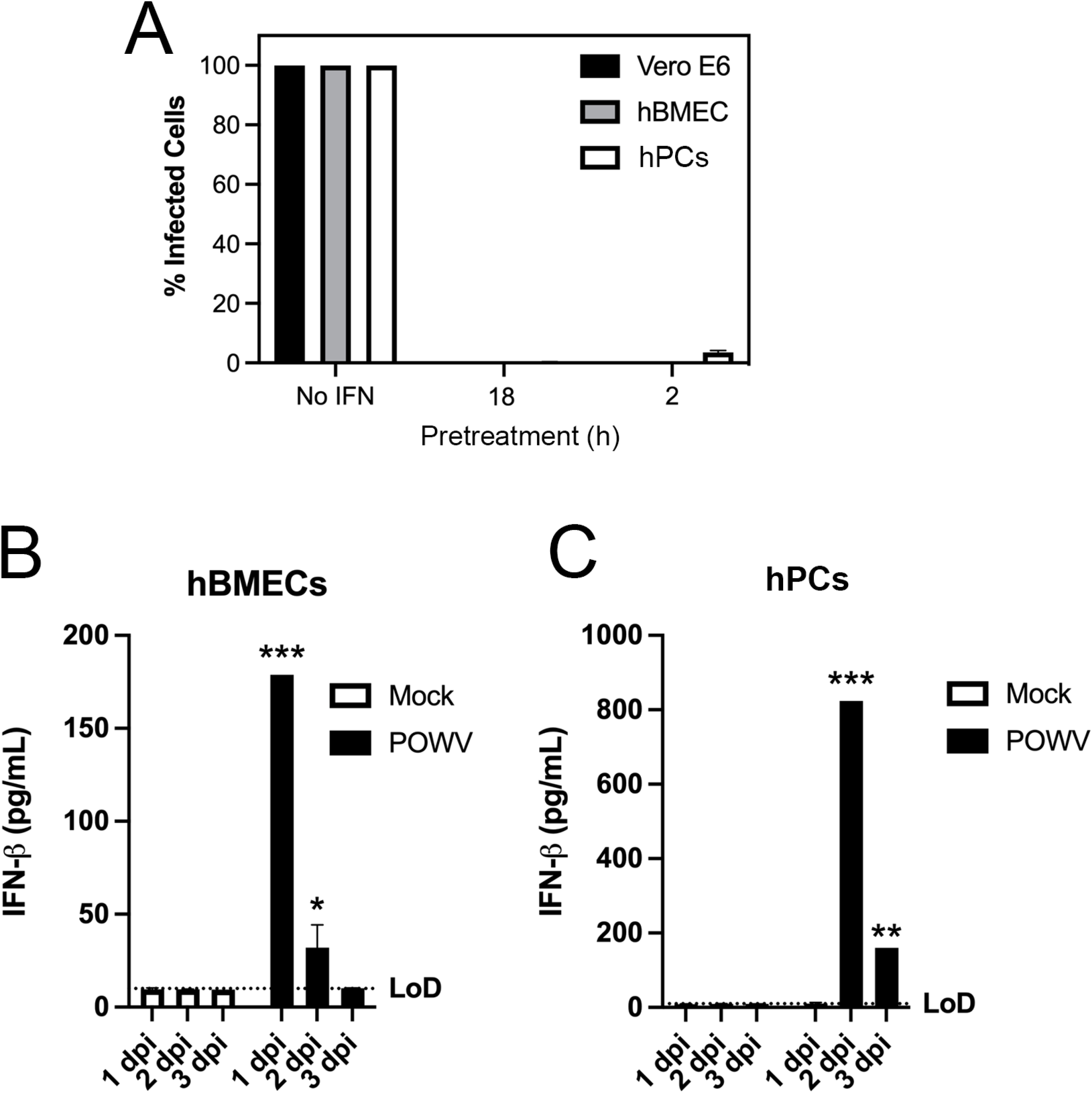
POWV-LI9 is susceptible to exogenous interferon addition. (A) Vero E6, hBMECs and human brain pericytes (hPCs) were pretreated with IFN-α (1,000 U/ml) for 2 or 18 h and before POWV infection (MOI, 5). One day post-infection cells were methanol fixed, immunoperoxidase stained using anti-POWV HMAF (1:10,000) and the number of infected cells quantitated and compared untreated controls. (B) hBMECs and (C) hPCs were mock or POWV-LI9 infected (MOI 5) and 1-3 dpi supernatants of cells were assayed for secreted IFNβ by ELISA (R&D systems).

### POWV Infected hBMECs and Pericytes Elicit Immune-Enhancing Responses

hBMECs and basolateral pericytes form a functional unit that determine the integrity of the BBB through synergistic responses that regulate vascular permeability, cell proliferation, and immune cell transit across the endothelium^(30, 31)^. To evaluate transcriptional changes of POWV infected hBMECs, we WGA treated POWV to synchronously infect 85-90% of cells 1 dpi and compared transcriptional responses to WGA treated, mock infected, hBMECs. RNA from control and POWV infected hBMECs were analyzed by Affymetrix arrays 1-3 dpi. POWV infection of hBMECs resulted in the dramatic induction of chemokines (CCL5, CXCL10, CCL20, CXCL11), IFNβ, IFN stimulated genes (ISGs: RSAD2, OASL, IFIT2/3, MX2, ISG15), IFN response factors (IRFs-1/2/7), Transcription factors (ATF3 and EGR1), and coagulation factor B over mock infected hBMEC controls (Figure 6A). Selected POWV directed hBMEC transcriptional changes are presented in Figure 5A, and complete data is available in the NCBI GEO database (GSE176251). POWV induction of high levels of IFNβ, CCL5, and ISG15 in hBMECs was verified by qRT-PCR (Figure 6B).

**Figure 6.**
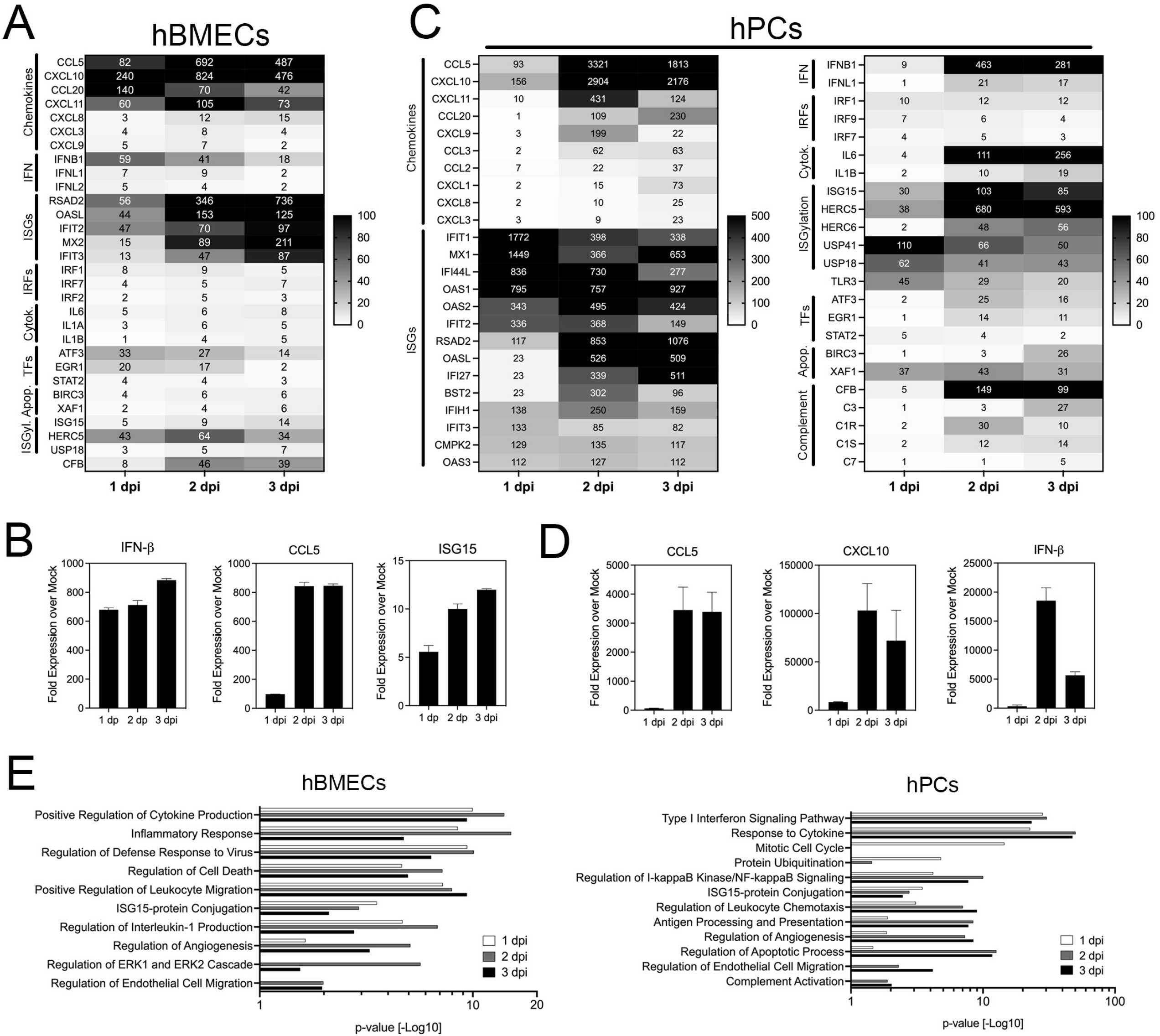
Global transcriptional changes in POWV-LI9-infected hBMECs and Pericytes. (A) POWV-LI9 was pretreated with WGA (1 μg/mL) for 5 min. Control mock WGA treated media (1 μg/mL), or WGA treated POWV was adsorbed to hBMECs for 1 hour, and subsequently cells were PBS washed resupplemented with EBM2-5% FBS media and grown for 1-3 days. Total cellular RNA from mock-WGA-treated and WGA-treated POWV infected hBMECs was harvested in RLT buffer, purified, DNAse treated and subjected to Affymetrix array analysis. Log_2_-induced genes from POWV-infected samples were compared to respective mock-infected time points, generating a heatmap with 0-to 100-fold induction scale, where induced genes above 100-fold were in black. Genes were separated into functional categories as chemokines, interferons (IFN), ISGs, IRFs, cytokines (Cytok.), transcriptional factors (TFs), apoptosis-related genes (Apop.), and ISGylation (ISGyl.) genes. (B) Selected genes from the Affymetrix array analysis were confirmed by qRT-PCR. (C) Total RNA from mock- and POWV-infected hPCs was isolated at 1-3 dpi and submitted to Affymetrix array analysis as in A. Log_2_-induced genes from POWV-infected samples were compared against its respective mock-infected timepoint, generating a heatmap with 0-to 500-fold (left graph) or 100-fold (right graph) induction scale, where induced genes above 500- or 100-fold were represented as black color. Genes were separated in functional categories as chemokines, interferons (IFN), ISGs, IRFs, cytokines (Cytok.), transcriptional factors (TFs), apoptosis-related (Apop.), ISGylation-related (ISGyl.), and complement related genes. (D) Selected genes induced by POWV infection of pericytes by Affymetrix array were confirmed by qRT-PCR. (E) Gene Ontology (GO) enrichment analysis was conducted by querying the upregulated genes from infected hBMECs or hPCs 1, 2, and 3 dpi using STRING v11^(32)^. GO terms from selected biological processes were presented.

We also analyzed human brain pericyte transcriptional responses following synchronous POWV-LI9 infection (MOI 5) compared to mock-infected pericytes (NCBI GEO accession GSE176252). ISGs were the most highly induced responses observed in POWV infected pericytes 1 dpi, with 100-1700 fold increases over a wide array of ISGs (Figure 6C). Despite high ISG responses, IFNβ was only induced 9 fold 1 dpi, but increased dramatically to 463-fold by 2 dpi and 281-fold 3 dpi. Chemokines CCL5 and CXCl10 were moderately induced 93-156 fold 1 dpi but were induced 1800-3300 fold 2-3 dpi (Figure 6C). Complement-activating factors (CFB, C3, C1S, C1R, C7), IL6 (111-256-fold 2-3 dpi) and IL1β (10-19-fold) were highly induced by POWV infection of pericytes 2-3 dpi. Despite high level IFNβ and ISG responses, POWV productively replicated and spread in pericytes 1-3 dpi, with persistent infection apparent >8 dpi (Figure 3D,E). The induction of selected chemokines, cytokines and ISGs induced by POWV infection was validated by qRT-PCR (Figure 6D). Our findings indicate that POWV infection of human pericytes transcriptionally induces chemotactic cytokine and complement responses that are known to enhance inflammation and immune cell recruitment to the endothelium. How POWV spreads and persists in pericytes in the presence of high level IFN and ISG responses at later times after infection remains to be investigated.

Pathway gene ontology (GO) analysis^(32)^ of hBMECs and pericyte transcriptomes showed similar GO enriched responses related to cytokine production, leukocyte migration, type I IFN signaling, ISG15-protein conjugation, cell death and apoptosis regulation, angiogenesis, and EC migration (Figure 6E). Regulation of prosurvival ERK1/2 signaling, that was recently shown to be required for ZIKV persistence in hBMECs^(33)^, was specifically associated with POWV induced hBMEC transcriptomes 2-3 dpi (Figure 6E, left panel). ERK1/2 pathway activation is consistent with CCR3/CCR5 activation by high levels of CCL5 induced by POWV-infected hBMECs and pericytes^(33)^. Pericyte specific responses included cell cycle, NF-κB signaling, antigen processing and presentation, and complement activation 2-3 dpi (Figure 5E, right panel).

Collectively, chemokine responses elicited by POWV infected hBMECs appear to be synergistic with robust chemokine and IFN responses induced by POWV-LI9 infected pericytes. These findings indicate that POWV infection of pericytes is likely to enhance immune cell recruitment to hBMECs that may contribute to immune enhanced pathogenesis. In contrast to IFN secretion 1 dpi by hBMECs, POWV infected pericytes inhibit IFN secretion 1 dpi, yet both cell types induce late IFN and ISG responses that fail to prevent POWV persistence. These findings suggest that POWV differentially regulates early IFN responses in pericytes and hBMECs, and suggest that at later times a subset of POWV infected hBMECs and pericytes may become resistant to IFN and ISG responses.

## Discussion

POWV is transmitted to humans in tick saliva during a short ~15 minute tick bite^(34)^. POWVs cause a severe neurovirulent disease that is 10% fatal and results in long term neurologic deficits in 50% of patients^(8)^. The rapid transmission of POWV is unlike Lyme disease, which requires long term tick attachment and where antibiotics are available to clear the infection. For POWV, there are no therapeutic treatments or approved vaccines to prevent neurovirulent infection and disease. POWV carrying ticks and human POWV infections have increased in the Northeastern (N.E.) U.S.^(22, 35, 36)^. Climate change is expected to exacerbate future POWV incidence by expanding the range and density of *I. scapularis* and mammalian disease reservoirs in addition to increasing tick survival and seasonal activity^(37)^. A recent study found a 2% prevalence of POWV in *Ixodes scapularis* ticks on Long Island, NY^(6)^, and the spread of POWV and the severity of disease rationalized evaluating POWVs circulating in this densely populated area. In this report, we isolated POWVs from *I. scapularis* ticks on LI and demonstrate that POWVs (LI9, LI41 and LB) infect human brain microvascular endothelial cells and pericytes that form a neurovascular complex that spans the BBB. Our findings suggest the potential for persistently infected hBMECs and pericytes to be reservoirs of POWV spread, and suggest a mechanism for POWVs to cross the BBB and enter neuronal compartments.

We isolated POWVs (LI9 and LI41) from 3/44 PCR positive LI tick pools by directly inoculating tick homogenates into VeroE6 cells. Cloning and sequencing of the POWV-LI9 tissue culture isolate revealed it to be a phylogenetically distinct lineage II POWV strain present in LI *Ixodes scapularis* ticks. The POWV-LI9 isolate is 99% identical to POWV genomes collected in Rhode Island and Maine (Figure 2A)^(20–22)^, and shares 95% amino acid identity with the prototypic 1958 lineage I POWV-LB strain (Figure 2B)^(20–22)^. However, other than sequencing, the isolation and tissue culture analysis of NE POWV strains has yet to be reported.

We isolated POWV-LI9 and POWV-LI41 by directly infecting VeroE6 cells with tick homogenates and observed large POWV infected cell foci in liquid media 7 dpi. In this setting, viruses are normally dispersed in cell supernatants and spread to single cells throughout the monolayer. The observation that POWV-LI9 and -LI4 form foci indicates that these POWVs spread cell to cell. This is novel and not previously evaluated or reported for any POWV^(38)^. In subsequent POWV-LI9 passages, focal spread observed in passage 1 and 2, changed to a more uniform infection of VeroE6 cells. Whether cell to cell spread is a fundamental feature of POWVs directly inoculated into mammalian cells has yet to be considered and remains to be determined as there are no studies of POWV spread following direct tissue culture isolation from ticks. Prototypic POWV-LB and POWV-SP strains were isolated by intracranial inoculation of mice prior to extensive passage in Vero cells^(1, 8, 14, 15)^. Prior studies of POWV-LB infected VeroE6 cells did not report infected cell foci, but also lack analysis of POWV spread in infected cells. We found that POWV-LB uniformly infected VeroE6 cells with a few grouped infected cells (Supp Fig 2), but no large infected cell foci indicative of cell to cell spread.

Initial cell to cell spread of POWV strains remains an enigma that requires analysis of more POWVs by direct isolation in tissue culture, and the analysis of potential POWV genetic changes during isolation and early passage. These findings also raise the question of whether POWVs are initially restricted to cell to cell spread following a tick bite, and whether this contributes to the long prodrome of POWV disease. These findings are fundamental to initial POWV spread and may need to be considered in studies of POWV spread and neuropathogenesis in murine models that are uniformly based on POWVs that were first passaged in mouse brain.

The ability of POWV-LI9 and -LI41 strains to form infected foci are new and novel observations, as there are no reports of cell to cell spread for other mosquito or tick-borne flaviviruses^(38)^. Focal cell to cell spread has been shown for several viruses including Herpesvirus, rhabdovirus, vaccinia virus, HIV-1 and the Flavivirus HCV^(38–44)^. Neurovirulent measles virus forms infectious foci in primary human airway epithelial cells, with viral spread dependent on intercellular adherens junction localized pores^(42)^. Similar to POWV, measles virus foci form without cytopathology, cell to cell fusion or syncytia formation^(42)^. Additional experiments that differentiate POWV cell to cell spread, basolateral release or spread via intercellular membrane pores are required to understand how POWV-LI9 forms foci, and to define mechanisms of POWV spread from infected hBMECs to the CNS.

How POWVs enter the CNS remains unknown and there is no information on cellular targets of POWV infection, or the ability of POWVs to infect and replicate in cells of the BBB. The BBB is a highly selective semipermeable barrier that separates the circulating blood from the CNS and prevents pathogens from entering the brain^(45)^. The BBB is composed of brain microvascular endothelial cells, pericytes, astrocytes and microglia^(31, 45, 46)^. Pericytes form intimate basolateral contacts with both BMECs and astrocytes, forming a neurovascular unit that regulates functions of luminal endothelial cells and abluminal pericytes and astrocytes within the neuronal compartment^(25, 47)^.

There are no prior studies of POWV infection of hBMECs and pericytes. We found that POWV-LI9, -LI41 and POWV-LB infect hBMECs resulting in single infected cells with little cell to cell spread after initial infection. This differs from POWV-LI9 infection of VeroE6 cells and it remains to be determined if these differences are the result of hBMEC receptors or IFN restricted spread that is absent in VeroE6 cells. POWVs productively infected hBMECs without apparent cell lysis or permeabilizing effects (Figure 4C). However, a comparison of POWV titers from apical and basolateral sides of polarized VeroE6 or hBMECs in Transwell plates demonstrated that POWV-LI9 is preferentially released from the basolateral side (3-12-fold) (Figure 4D, E). Similar analysis of POWV-LB found both apical and basolateral spread from infected hBMECs with a 3-fold increase in basolateral spread (Supp Fig 4B). Only a few studies demonstrate preferential basolateral viral spread^(48–50)^. Vesicular stomatitis virus (VSV) and Marburg virus (MARV) bud from the basolateral side of polarized MDCK cells^(48–50)^, and arenaviruses are basolaterally released from airway epithelial cells^(51)^. Basolateral spread of POWVs suggests a mechanism for POWVs to cross the BBB and spread abluminally to neuronal compartments.

In the brain, pericytes form a neurovascular complex with hBMECs spanning the BBB and contacting astrocyte end feet, microglia, and neurons^(30, 31, 52)^. POWV infection of hBMECs prompted us to analyze whether pericytes are infected and whether hBMECs convey POWV infection to pericytes. We found that POWV-LI9, LI41 and POWV-LB highly infect primary human brain pericytes (Figure 3D,E; Supp Fig. 4A,C,E). In a BBB-like in vitro model POWV-LI9 infected hBMECs also basolaterally transmitted POWV to pericytes in the lower chamber (Figure 4F). Our findings demonstrate that POWVs productively and persistently infect hBMECs and pericytes, and suggest hBMEC-pericyte complexes as potential POWV reservoirs and conduits for viral spread to neuronal compartments.

Neurotropic flaviviruses including ZIKV, JEV, WNV and TBEV are reportedly transmitted across polarized ECs without permeabilizing or lysing cells^(16, 18, 53, 54)^. In TBEV infected mice, BBB permeability was not required for viral entry into the CNS, and permeability was only observed at late stages of infection, and after high viral loads and inflammatory damage was evident in the brain^(17)^. In innate immune compromised mice, neurotropic flaviviruses invade the CNS during early viremic stages^(55)^ with infection of the BBB one of the most likely routes of neuronal invasion^(45)^.

Many murine POWV pathogenesis models use POWV-LB and -SP strains that were first passaged in mouse brains^(14, 56)^, and IFNAR antibody blockades to promote lethal outcomes^(57, 58)^. Initial inoculation of POWV-LI9 strain in WT C57BL/6 mice resulted in murine seroconversion and antibodies that neutralized both POWV-LI9 and POWV-LB (Supp. Fig. 3A,B). It remains to be determined if POWV-LI9 and −41 strains are neurovirulent in mice. However, analysis of POWV-LI9 viremia, CNS spread, pathology and neurovirulence in mice, at early or late passage, is beyond the scope of this initial study of newly isolated POWV strains circulating on LI.

POWV-LI9 persistence in hBMECs and pericytes is consistent with the delayed onset of neurologic disease in POWV patients and the long-term pathology of POWV infection^(4, 8, 59)^. However, POWV persistence is very different from the persistent infection of hBMECs by ZIKV^(18, 33)^. ZIKV replicates and spreads persistently in hBMECs without permeabilizing or lysing cells, and persistence continues through serial cellular passage^(18)^. Recent findings indicate that ZIKV persistence requires the suppression of IFN secretion and the high level secretion of CCL5 that acts as an autocrine activator of ERK1/2 survival signaling responses^(33)^.

In contrast to ZIKV, POWV infection of hBMECs does not result in the constitutive spread of POWV throughout the monolayer. ZIKV, DENV and POWV are inhibited by pretreating cells with Type I IFN^(18, 60)^. However, ZIKV bypasses this restriction by blocking IFNβ secretion from infected hBMECs, while DENV is cleared from infected ECs by IFNβ and fails to establish a persistent infection^(18, 60)^. Although, POWV spread is restricted by secreted IFNβ, a subset of infected hBMECs remain persistently infected, viable and POWV antigen positive. Curiously, POWV inhibits pericyte secretion of IFNβ 1 dpi, permitting early viral spread within pericytes. This suggests novel POWV regulation of early IFN responses in pericytes and suggests that POWV resistance to IFN secreted 2-3 dpi permits POWV to persistently infect cells. It remains unknown whether POWV proteins, generated early in infection, permit POWV persistence or resistance to IFN/ISG responses hBMECs or pericytes or how secreted IFNβ levels 3 dpi are restricted, despite highly induced IFNβ transcripts. Collectively these findings indicate that POWV persists in hBMECs and pericytes without a total blockade of IFNβ secretion observed during ZIKV infection^(18)^.

BMECs basolaterally secrete PDGF which recruits and stabilizes pericyte binding to BMECs and reciprocally, pericytes secrete factors that act on BMEC receptors which regulate barrier integrity, angiogenesis, and immune cell recruitment^(25, 47)^. Pericytes regulate the BBB, and targeting PDGFs or pericyte-BMEC interactions using imatinib or nintedanib has been suggested as a potential therapeutic approach to control damage of this neurovascular complex^(25, 47, 61)^. Our analysis of POWV-LI9 directed changes in hBMEC and pericyte transcriptional responses included finding the induction of several pro-survival genes including EGR1, ATF3 and BIRC3^(62–64)^, and chemokine CCL5, a cell survival response required for ZIKV persistence in hBMECs^(33)^. Highly induced CCL5 activates ERK1/2 cell survival pathways in ECs^(33)^ that may regulate anti-apoptotic and proliferative responses^(65)^ and contribute to the viability of POWV infected hBMECs and pericytes.

Several chemokines were upregulated in both cell types, however, CCL5 and CXCL10 were the most highly induced, especially by POWV infected pericytes (Figure 6). CXCL10 is prominently upregulated in the CSF of human patients and mouse models during TBEV and WNV infections^(66–70)^ and higher CXCL10 expression in the brain is correlated with more severe TBEV disease in mice^(71)^. CCL5 and CXCL10 also play an important role in recruiting activated T cells into sites of tissue inflammation^(72)^ and these chemokines are implicated in both viral clearance and the neuropathology of neurotropic flavivirus infections^(73)^. CCL5 is produced by several cell types that impact inflammation in the CNS, including hBMECs, pericytes, astrocytes, microglia, and neurons^(73, 74)^. Increased CCL5 has been detected in the CSF, but not in the serum, of TBEV patients and TBEV-infected mice^(68, 75)^. During TBEV infections, CCL5-directed neuroinflammation may contribute to brain pathology as treatment of mice with CCL5 antagonists prolonged survival and reduced brain inflammation after TBEV infection^(76)^. Similarly, the CCL5 receptor antagonist, Maraviroc, reduced JEV-induced CNS inflammation and increased the survival of JEV infected mice^(77)^. CCR5 knockout mice were also protected from fatal WNV or JEV encephalitis^(70, 78)^. ZIKV persistence in hBMECs was recently shown to require high level CCL5 secretion to support prosurvival responses, and CCL5 receptor inhibitors, that prevent ZIKV persistence in hBMECs, are suggested as therapeutics that clear ZIKV infections^(33)^.

We were surprised to find that complement related factors CFB (149-fold), C3 (27-fold), C1r (30-fold), and C1s (14-fold) were highly induced by POWV infected pericytes. Complement proteins in the brain can be induced by astrocytes and microglia in response to injury^(79)^, but little is known about brain pericytes actively producing complement proteins. WNV and HIV-1 induce complement activation within the CNS^(80, 81)^, and elevated C1q levels were found in the CSF of WNV and TBEV infected patients^(82)^, and suggested to contribute to neurocognitive impairment in recovering patients^(83)^. Our findings suggest that POWV-infected pericytes might enhance complement activation and C3a and C5a production, which contribute to immune cell chemotaxis, and T cell priming responses involved in both viral clearance and POWV neuropathology.

In this study, we demonstrate that POWVs productively infect and are basolaterally released from brain endothelial cells, suggesting a potential mechanism by which POWVs infect pericytes and cross neurovascular complexes within the BBB. A subset of hBMECs and pericytes are persistently infected by POWV for 8-30 days providing a novel location for the replication and spread of a neurovirulent virus to the CNS. The abluminal location of pericytes in association with BBB hBMEC makes them ideally suited to convey POWV to the CNS and recruit immune cells to the BBB that contribute to neuronal immunopathogenesis. The ability of POWV to persist in hBMECs and pericytes also suggests that these cellular niches are potential therapeutic targets for preventing POWV neurovirulence.

## Material and Methods

### Cells

VeroE6 cells (ATCC CRL 1586) were grown in DMEM (Dulbecco’s modified Eagle’s medium) supplemented with 5% fetal bovine serum (FBS) and penicillin (100 μg/ml), streptomycin sulfate (100 μg/ml), and amphotericin B (50 μg/ml; Mediatech) at 37°C and 5% CO_2_. hBMECs (passages 4 to 10) were purchased from Cell Biologics (H-6023) and grown in EC basal medium-2 MV (EBM-2 MV; Lonza) supplemented with EGM-2 MV SingleQuots (Lonza) and incubated at 37°C and 5% CO_2_. Human brain vascular pericytes (hPCs - passage 1) were purchased from ScienCell (#1200) and grown in pericyte medium (PM, #1201) supplemented with pericyte growth supplement (#1252) and incubated at 37°C and 5% CO_2_.

### POWV-LI9 Isolation from *Ixodes scapularis* Ticks

Ticks were collected during the fall season peak (October to November) throughout Suffolk County, New York, in 2020. Questing ticks were collected from vegetation along trails by flagging a 1-m^2^ cotton flannel fabric attached on both ends to a wooden pole between 10:00 and 14:00 h during sunny days. 438 adults *Ixodes scapularis* ticks were identified morphologically on a dissecting microscope for taxonomic keys. Pools of 10 ticks were homogenized using an eppendorf tube mortar and pestle in 500 μL of phosphate-buffered saline (PBS) and debris removed by centrifuging for 5 min at 12,000 x g.

### qRT-PCR for tick screening

Tick pool homogenates were tested for POWV RNA using a multiplex one-step reverse transcription PCR (qRT-PCR) using a forward primer (GTGATGTGGCAGCGCACC), a reverse primer (CTGCGTCGGGAGCGACCA) and a probe (CCTACTGCGGCAGCACACACAGTG) (6). The final reaction mixture contained 5 μl of template and 20 μl of master mixture. The master mix contained 0.2 μM of forward primer, 0.2 μM of reverse primer, 0.1 μM of probe, 1.25 μl of RNA UltraSense enzyme mix, and 5 μl RNA UltraSense 5× reaction mix. The reverse transcription step was performed at 55°C for 15 min, followed by incubation at 95°C for 10 min. The PCR consisted of 40 cycles at 95°C for 15 s and 60°C for 30 s. The qPCR was performed on a Bio-Rad C1000 Touch system with a CFX96 optical module (Bio-Rad) using the RNA UltraSense one-step quantitative RT-PCR system protocol (Invitrogen).

### Inoculation of Mammalian Cells

Tick pool homogenates were inoculated onto individual 24 well or 6-well plates of VeroE6 cells (150 μl/well) in DMEM (supplemented with 2% FBS; penicillin 100 μg/ml; streptomycin sulfate 100 μg/ml; and amphotericin B 50 μg/ml). Plates were transferred to the Stony Brook BSL3 facility for growth at 37°C and 5% CO_2_. At 7-10 dpi cell supernatants were harvested and cell monolayers were 100% methanol fixed and POWV infected cells detected by immunoperoxidase staining with anti-POWV HMAF sera (ATCC). Supernatants positive for POWV RNA by qRT-PCR and POWV antigen staining in infected cells and were inoculated into T25 flasks containing VeroE6 cells and propagated for 7 days in DMEM with 2% FBS.

### POWV-LI9 Infection and Immunoperoxidase staining

POWV-LI9 was adsorbed to ~60% confluent hBMEC or human pericyte monolayers for 1 h. Following adsorption, monolayers were washed with PBS and grown in supplemented EBM-2 MV or PM with 5% FBS, respectively. VeroE6 cells were similarly infected, washed, and supplemented with DMEM with 8% FBS. POWV-LI9 titers were determined by serial dilution and infection of VeroE6 cells, quantifying infected cell foci at 24 hpi by immunoperoxidase staining with anti-POWV hyperimmune mouse ascites fluid (HMAF; 1:5,000; ATCC), horseradish peroxidase (HRP)-labeled anti-mouse IgG (1:2,000; KPL-074-1806), and 3-amino-9-ethylcarbazole staining^(60)^. IFN-α inhibition assay, medium was supplemented with 1,000 U/ml IFN-α (Sigma-Aldrich) at indicated times and cells incubated at 37°C and 5% CO_2_.

### POWV-LI9 Cloning and Sequencing

Total RNA was purified using RNeasy (Qiagen) from early passage POWV-infected Vero E6 cells (passage 3). cDNA synthesis was performed using a Transcriptor first-strand cDNA synthesis kit (Roche) using random hexamers as primers (25°C for10 min, 50°C for 60 min, and 90°C for 5 min). POWV genome was amplified using primer pairs to regions located at near the junctions between capsid (forward: AGGAGAACAAGAGCTGGGAGTGGTC) and envelope (reverse: TGCTCCGACTCCCATTGTCATCAT); NS1 (forward: GACTATGGATGTGCAGTTGATCC) and NS2B (reverse: TCTGCGTGCTGATGAGAAGA); NS3 (forward: ACTGACCTTGTATTCTCAGGG) and NS4B (reverse: GTGCATGAGTTCAACCGTTG); and NS5 (forward: CTAGAAGGGGTGGAGCAGAGG; reverse: TTAGATTATTGAGCTCTCTAGCTTG). 5’UTR was obtained using the Template Switching RT Enzyme Mix (NEB #M0466) following the manufacturer’s protocol. 3’ UTR was obtained using the reverse primer (GCGGGTGTTTTTCCGAGTCACACAC). PCR fragments were obtained using Phusion polymerase (NEB M0530L) with 37 cycles at 98°C for 30s, 68°C for 30s, and 72°C using 30s per kb. PCR fragments were gel purified (NEB T1020L) and cloned into pMiniT 2.0 vector (NEB E1202S). POWV-LI9 genome was sequenced using vector and internal primers, assembling both strands at least twice and submitted to GenBank (MZ576219).

### Phylogenetic analysis

A phylogenetic tree of POWV sequences was generated using the neighbor-joining method^(84)^ with a maximum composite likelihood model and bootstrap test of phylogeny with 100 replications^(85)^ in MEGA version X (86). Phylogenetic analysis was performed with the POWV-LI9 sequence and reference full genome sequences representing POWV lineages I and II, and TBEV as outgroup, obtained from NCBI, as follows: Lineage I (LB L06436, LEIV-5530 KT224351, Ternay HQ231415, Pow-24 MG652438, Ulysses HQ231414, Spassk-9 EU770575, Nadezdinsk-1991 EU670438, LEIV-3070Prm KT224350, POWPa06 EU543649, POWANY64-7062 HM440563), Lineage II (RTS81 MG647779, RTS82 MG647780, RTS84 MG647783, RTS92 MG647781, RTS96 MG647782, NSF001 HM440559, MeW17-228 MK309362, P0375 KU886216, LI-1 KJ746872, DTVMeC17 MK104144, CTB30 AF311056, DTVWi99 HM440558, DTVWiA08 HM440560, DTVWiB08 HM440561, DTVWiC08 HM440562), and TBEV (NC_001672).

### Viability assay

PI (Calbiochem) and calcein-AM (Invitrogen) uptake were used to evaluate Vero E6 and hBMEC viability as previously described^(18)^. Briefly, POWV-infected (MOI, 5) or mock-infected cells were seeded into 24-well plates and co-stained by the membrane-permeable dye calcein-AM (3 μM; green fluorescence in live cells), and 2.5 μM propidium iodide (red fluorescent DNA stain to detect dead cells) 3 dpi. Images of calcein-AM-positive versus PI-positive cells were resolved using an Olympus IX51 microscope and Olympus DP71 camera.

### Confocal Immunofluorescence

POWV-LI9 (MOI, 5) was adsorbed to 80% confluent VeroE6 monolayers cultured in Lab-Tek II chambers (Nunc), washed with PBS, and grown in DMEM with 8% FBS during 48h. Cells were washed with PBS, fixed for 10 min with 4% paraformaldehyde-PBS, and permeabilized with 0.1% Triton X-100 in PBS for 10 min. Cells were blocked using 5% bovine serum albumin in PBS for 2 h and incubated with anti-POWV HMAF (1:10,000) and anti-ZO1 rabbit monoclonal antibody (1:100) in blocking solution for 18 h at 4°C. Cells were washed and incubated for 2 h with Alexa 488-conjugated goat anti-mouse IgG antibody (Invitrogen) and with Alexa 546-conjugated goat anti-rabbit IgG antibody (Invitrogen) diluted 1:400 in blocking solution at room temperature. hBMECs were subsequently incubated with 5 μM 4,6-diamidino-2-phenylindole (DAPI; Sigma) for 5 min at room temperature. Slides were mounted using ProLong Antifade (Thermo Fisher) solution and observed using a Zeiss LSM 510 META/NLO confocal microscope.

### Affymetrix gene array analysis

Primary hBMECs or human pericytes (passages 4 to 10) were synchronously infected with POWV-LI9 (MOI, 5) or mock infected. For hBMECs only, POWV-LI9 or equivalent media were incubated with WGA (1 μg/ml) for 5 minutes prior to adsorbing POWV to cells for 1 hour, or mock infecting cells with equivalent amounts of WGA as controls. Following adsorption, cells were washed with PBS and media was replaced. Total RNA was purified 1-3 dpi from mock- or POWV-infected hBMECs or hPCs using RLT lysis buffer and RNeasy columns (Qiagen). Purified RNA was quantitated, and transcriptional responses detected on Affymetrix Clariom-S chip arrays in the Stony Brook Genomics Core Facility. POWV-infected cell transcriptional responses were compared to those in mock-infected cells harvested at each time point and fold changes in POWV versus control transcripts were analyzed using Affymetrix TAC software. Data obtained from these studies were submitted to the NCBI Gene Expression Omnibus database (hBMECs: GEO GSE176251; hPCs: GEO GSE176252).

### qRT-PCR analysis

Quantitative real-time PCR was performed on purified cellular RNAs derived from mock- or POWV-infected hBMECs/hPCs as described above. cDNA synthesis was performed using a Transcriptor first strand cDNA synthesis kit (Roche) using random hexamers as primers (25°C for 10 min, 50°C for 60 min, and 90°C for 5 min). qRT-PCR primers for specific genes were designed according to the NCBI gene database with 60°C annealing profiles (provided by Operon). Genes were analyzed using PerfeCTa SYBR green SuperMix (Quanta Biosciences) on a Bio-Rad C1000 Touch system with a CFX96 optical module (Bio-Rad). Responses were normalized to internal glyceraldehyde-3-phosphate dehydrogenase (GAPDH) mRNA levels, and the fold induction was calculated using the 2^-ΔΔCT^ method for differences between mock- and POWV-infected RNA levels at each time point.

### Trans Endothelial/Epithelial Electrical Resistance (TEER)

hBMECs or VeroE6 were plated on Costar Transwell inserts (3-μm pore size; Corning) at high density, and 7 days post seeding monolayers were analyzed for TEER (EVOM2; STX3; World Precision Instruments, Inc.). Confluent Transwell cultures were infected with POWV-LI9 (MOI, 5) or mock infected, and TEER values for POWV-infected versus mock-infected cells were compared at 1 to 3 dpi. For analysis of apical and basolateral release of POWV-LI9, hBMECs and VeroE6 were similarly infected 3 dpi prior to analyzing viral titers in apical and basolateral supernatants.

### POWV Transcytosis Assay

hBMECs were plated on Costar Transwell inserts (3-μm pore size; Corning) at high density, and 7 days post seeding hBMECs were mock- or POWV-infected in the upper chambers for 1 h, cells were washed with PBS, and the inserts were transferred to 24-well plates containing 70% confluent primary human brain vascular pericytes.

### ELISA

IFN-β in the supernatants of mock- and POWV-infected hBMECs or hPCs at 1, 2, and 3 dpi were measured using a DuoSet ELISA (R&D Systems). ELISA plates (Immunolon 2, U-bottom; Dynatech Laboratories) were coated with anti-IFN-β capture antibody according to the manufacturer. Viral supernatants were incubated with antibody coated plates (2 h), washed with PBS (0.1% Tween 20), and bound protein was detected with IFN-β specific antibodies conjugated to streptavidin-HRP and developed using tetramethylbenzidine. Secreted IFN-β in samples were compared to a purified IFN-β (R&D Systems) standard curve using a BioTek EL312e microplate reader (450 nm).

### Statistical analysis

Results shown in each figure were derived from two to three independent experiments with comparable findings; the data presented are means standard errors of the means (SEM), with the indicated P values of <0.01 and <0.001 considered significant. Two-way comparisons were performed two-tailed analysis of variance and an unpaired Student’s t test. All analyses were performed using GraphPad Prism software version 9.0.

## Data availability

Sequencing data from POWV-LI9 was submitted to GenBank (MZ576219). Microarray data obtained from these studies were submitted to the NCBI Gene Expression Omnibus database (hBMECs: GSE176251; hPCs: GSE176252).

## Acknowledgements

We thank Patrick Hearing, Carol Carter, Grace Himmler, and Nancy Reich-Marshall for helpful discussions and manuscript feedback; and Smruti Mishra and Luke Helminiak for animal monitoring and sample retrieval.

## References

1. Mc LD, Donohue WL. 1959. Powassan virus: isolation of virus from a fatal case of encephalitis. Can Med Assoc J 80:708–11.

2. Normandin E, Solomon IH, Zamirpour S, Lemieux J, Freije CA, Mukerji SS, Tomkins-Tinch C, Park D, Sabeti PC, Piantadosi A. 2020. Powassan Virus Neuropathology and Genomic Diversity in Patients With Fatal Encephalitis. Open Forum Infect Dis 7:ofaa392.

3. Robich RM, Cosenza DS, Elias SP, Henderson EF, Lubelczyk CB, Welch M, Smith RP. 2019. Prevalence and Genetic Characterization of Deer Tick Virus (Powassan Virus, Lineage II) in Ixodes scapularis Ticks Collected in Maine. Am J Trop Med Hyg 101:467–471.

4. Hermance ME, Thangamani S. 2017. Powassan Virus: An Emerging Arbovirus of Public Health Concern in North America. Vector Borne Zoonotic Dis 17:453–462.

5. El Khoury MY, Camargo JF, White JL, Backenson BP, Dupuis AP, 2nd, Escuyer KL, Kramer L, St George K, Chatterjee D, Prusinski M, Wormser GP, Wong SJ. 2013. Potential role of deer tick virus in Powassan encephalitis cases in Lyme disease-endemic areas of New York, U.S.A. Emerg Infect Dis 19:1926–33.

6. Sanchez-Vicente S, Tagliafierro T, Coleman JL, Benach JL, Tokarz R. 2019. Polymicrobial Nature of Tick-Borne Diseases. mBio 10.

7. Fatmi SS, Zehra R, Carpenter DO. 2017. Powassan Virus-A New Reemerging Tick-Borne Disease. Front Public Health 5:342.

8. Kemenesi G, Banyai K. 2019. Tick-Borne Flaviviruses, with a Focus on Powassan Virus. Clin Microbiol Rev 32.

9. Khan M, Beckham JD, Piquet AL, Tyler KL, Pastula DM. 2019. An Overview of Powassan Virus Disease. Neurohospitalist 9:181–182.

10. Feder HM, Telford S, Goethert HK, Wormser GP. 2020. Powassan Virus Encephalitis Following Brief Attachment of Connecticut Deer Ticks. Clin Infect Dis doi:10.1093/cid/ciaa1183.

11. Tavakoli NP, Wang H, Dupuis M, Hull R, Ebel GD, Gilmore EJ, Faust PL. 2009. Fatal case of deer tick virus encephalitis. N Engl J Med 360:2099–107.

12. Donoso Mantke O, Schadler R, Niedrig M. 2008. A survey on cases of tick-borne encephalitis in European countries. Euro Surveill 13.

13. Ebel GD. 2010. Update on Powassan virus: emergence of a North American tick-borne flavivirus. Annu Rev Entomol 55:95–110.

14. Ebel GD, Foppa I, Spielman A, Telford SR, 2nd. 1999. A focus of deer tick virus transmission in the northcentral United States. Emerg Infect Dis 5:570–4.

15. Corrin T, Greig J, Harding S, Young I, Mascarenhas M, Waddell LA. 2018. Powassan virus, a scoping review of the global evidence. Zoonoses Public Health 65:595–624.

16. Palus M, Vancova M, Sirmarova J, Elsterova J, Perner J, Ruzek D. 2017. Tick-borne encephalitis virus infects human brain microvascular endothelial cells without compromising blood-brain barrier integrity. Virology 507:110–122.

17. Ruzek D, Salat J, Singh SK, Kopecky J. 2011. Breakdown of the blood-brain barrier during tick-borne encephalitis in mice is not dependent on CD8+ T-cells. PLoS One 6:e20472.

18. Mladinich MC, Schwedes J, Mackow ER. 2017. Zika Virus Persistently Infects and Is Basolaterally Released from Primary Human Brain Microvascular Endothelial Cells. MBio 8.

19. Rosales-Munar A, Alvarez-Diaz DA, Laiton-Donato K, Pelaez-Carvajal D, Usme-Ciro JA. 2020. Efficient Method for Molecular Characterization of the 5’ and 3’ Ends of the Dengue Virus Genome. Viruses 12.

20. Pesko KN, Torres-Perez F, Hjelle BL, Ebel GD. 2010. Molecular epidemiology of Powassan virus in North America. J Gen Virol 91:2698–705.

21. Anderson JF, Armstrong PM. 2012. Prevalence and genetic characterization of Powassan virus strains infecting Ixodes scapularis in Connecticut. Am J Trop Med Hyg 87:754–9.

22. Tokarz R, Williams SH, Sameroff S, Sanchez Leon M, Jain K, Lipkin WI. 2014. Virome analysis of Amblyomma americanum, Dermacentor variabilis, and Ixodes scapularis ticks reveals novel highly divergent vertebrate and invertebrate viruses. J Virol 88:11480–92.

23. Attwell D, Mishra A, Hall CN, O’Farrell FM, Dalkara T. 2016. What is a pericyte? J Cereb Blood Flow Metab 36:451–5.

24. He Y, Yao Y, Tsirka SE, Cao Y. 2014. Cell-culture models of the blood-brain barrier. Stroke 45:2514–26.

25. Liu Q, Yang Y, Fan X. 2020. Microvascular pericytes in brain-associated vascular disease. Biomed Pharmacother 121:109633.

26. Wang S, Cao C, Chen Z, Bankaitis V, Tzima E, Sheibani N, Burridge K. 2012. Pericytes regulate vascular basement membrane remodeling and govern neutrophil extravasation during inflammation. PLoS One 7:e45499.

27. Banks WA, Broadwell RD. 1994. Blood to brain and brain to blood passage of native horseradish peroxidase, wheat germ agglutinin, and albumin: pharmacokinetic and morphological assessments. J Neurochem 62:2404–19.

28. Banks WA, Akerstrom V, Kastin AJ. 1998. Adsorptive endocytosis mediates the passage of HIV-1 across the blood-brain barrier: evidence for a post-internalization coreceptor. J Cell Sci 111 (Pt 4):533–40.

29. Banks WA, Kastin AJ. 1998. Characterization of lectin-mediated brain uptake of HIV-1 GP120. J Neurosci Res 54:522–9.

30. Rustenhoven J, Jansson D, Smyth LC, Dragunow M. 2017. Brain Pericytes As Mediators of Neuroinflammation. Trends Pharmacol Sci 38:291–304.

31. Winkler EA, Bell RD, Zlokovic BV. 2011. Central nervous system pericytes in health and disease. Nat Neurosci 14:1398–1405.

32. Szklarczyk D, Gable AL, Lyon D, Junge A, Wyder S, Huerta-Cepas J, Simonovic M, Doncheva NT, Morris JH, Bork P, Jensen LJ, Mering CV. 2019. STRING v11: protein-protein association networks with increased coverage, supporting functional discovery in genome-wide experimental datasets. Nucleic Acids Res 47:D607–D613.

33. Mladinich M, Conde JN, Schutt WR, Sohn SY, Mackow ER. 2021. Blockade of Autocrine CCL5 Responses Inhibits Zika Virus Persistence and Spread in Human Brain Microvascular Endothelial Cells. MBio 12:e01962–21.

34. Ebel GD, Kramer LD. 2004. Short report: duration of tick attachment required for transmission of powassan virus by deer ticks. Am J Trop Med Hyg 71:268–71.

35. Smith RP, Jr., Elias SP, Cavanaugh CE, Lubelczyk CB, Lacombe EH, Brancato J, Doyle H, Rand PW, Ebel GD, Krause PJ. 2019. Seroprevalence of Borrelia burgdorferi, B. miyamotoi, and Powassan Virus in Residents Bitten by Ixodes Ticks, Maine, USA. Emerg Infect Dis 25:804–807.

36. Yuan Q, Llanos-Soto SG, Gangloff-Kaufmann JL, Lampman JM, Frye MJ, Benedict MC, Tallmadge RL, Mitchell PK, Anderson RR, Cronk BD, Stanhope BJ, Jarvis AR, Lejeune M, Renshaw RW, Laverack M, Lamb EM, Goodman LB. 2020. Active surveillance of pathogens from ticks collected in New York State suburban parks and schoolyards. Zoonoses Public Health 67:684–696.

37. Bouchard C, Dibernardo A, Koffi J, Wood H, Leighton PA, Lindsay LR. 2019. N Increased risk of tick-borne diseases with climate and environmental changes. Can Commun Dis Rep 45:83–89.

38. Zhong P, Agosto LM, Munro JB, Mothes W. 2013. Cell-to-cell transmission of viruses. Curr Opin Virol 3:44–50.

39. Timpe JM, Stamataki Z, Jennings A, Hu K, Farquhar MJ, Harris HJ, Schwarz A, Desombere I, Roels GL, Balfe P, McKeating JA. 2008. Hepatitis C virus cell-cell transmission in hepatoma cells in the presence of neutralizing antibodies. Hepatology 47:17–24.

40. Xiao F, Fofana I, Heydmann L, Barth H, Soulier E, Habersetzer F, Doffoel M, Bukh J, Patel AH, Zeisel MB, Baumert TF. 2014. Hepatitis C virus cell-cell transmission and resistance to direct-acting antiviral agents. PLoS Pathog 10:e1004128.

41. Sattentau Q. 2008. Avoiding the void: cell-to-cell spread of human viruses. Nat Rev Microbiol 6:815–26.

42. Singh BK, Hornick AL, Krishnamurthy S, Locke AC, Mendoza CA, Mateo M, Miller-Hunt CL, Cattaneo R, Sinn PL. 2015. The Nectin-4/Afadin Protein Complex and Intercellular Membrane Pores Contribute to Rapid Spread of Measles Virus in Primary Human Airway Epithelia. J Virol 89:7089–96.

43. Schiffner T, Sattentau QJ, Duncan CJ. 2013. Cell-to-cell spread of HIV-1 and evasion of neutralizing antibodies. Vaccine 31:5789–97.

44. Farnsworth A, Johnson DC. 2006. Herpes simplex virus gE/gI must accumulate in the trans-Golgi network at early times and then redistribute to cell junctions to promote cell-cell spread. J Virol 80:3167–79.

45. Cain MD, Salimi H, Diamond MS, Klein RS. 2019. Mechanisms of Pathogen Invasion into the Central Nervous System. Neuron 103:771–783.

46. Liu WY, Wang ZB, Zhang LC, Wei X, Li L. 2012. Tight junction in blood-brain barrier: an overview of structure, regulation, and regulator substances. CNS Neurosci Ther 18:609–15.

47. Armulik A, Genove G, Mae M, Nisancioglu MH, Wallgard E, Niaudet C, He L, Norlin J, Lindblom P, Strittmatter K, Johansson BR, Betsholtz C. 2010. Pericytes regulate the blood-brain barrier. Nature 468:557–61.

48. Compton T, Ivanov IE, Gottlieb T, Rindler M, Adesnik M, Sabatini DD. 1989. A sorting signal for the basolateral delivery of the vesicular stomatitis virus (VSV) G protein lies in its luminal domain: analysis of the targeting of VSV G-influenza hemagglutinin chimeras. Proc Natl Acad Sci U S A 86:4112–6.

49. Kolesnikova L, Ryabchikova E, Shestopalov A, Becker S. 2007. Basolateral budding of Marburg virus: VP40 retargets viral glycoprotein GP to the basolateral surface. J Infect Dis 196 Suppl 2:S232–6.

50. Drokhlyansky E, Soh TK, Cepko CL. 2015. Preferential Budding of Vesicular Stomatitis Virus from the Basolateral Surface of Polarized Epithelial Cells Is Not Solely Directed by Matrix Protein or Glycoprotein. J Virol 89:11718–22.

51. Dylla DE, Michele DE, Campbell KP, McCray PB, Jr. 2008. Basolateral entry and release of New and Old World arenaviruses from human airway epithelia. J Virol 82:6034–8.

52. Stebbins MJ, Gastfriend BD, Canfield SG, Lee MS, Richards D, Faubion MG, Li WJ, Daneman R, Palecek SP, Shusta EV. 2019. Human pluripotent stem cell-derived brain pericyte-like cells induce blood-brain barrier properties. Sci Adv 5:eaau7375.

53. Hsieh JT, St John AL. 2020. Japanese encephalitis virus and its mechanisms of neuroinvasion. PLoS Pathog 16:e1008260.

54. Verma S, Lo Y, Chapagain M, Lum S, Kumar M, Gurjav U, Luo H, Nakatsuka A, Nerurkar VR. 2009. West Nile virus infection modulates human brain microvascular endothelial cells tight junction proteins and cell adhesion molecules: Transmigration across the in vitro blood-brain barrier. Virology 385:425–33.

55. Suen WW, Prow NA, Hall RA, Bielefeldt-Ohmann H. 2014. Mechanism of West Nile virus neuroinvasion: a critical appraisal. Viruses 6:2796–825.

56. Hermance ME, Hart CE, Esterly AT, Reynolds ES, Bhaskar JR, Thangamani S. 2020. Development of a small animal model for deer tick virus pathogenesis mimicking human clinical outcome. PLoS Negl Trop Dis 14:e0008359.

57. VanBlargan LA, Himansu S, Foreman BM, Ebel GD, Pierson TC, Diamond MS. 2018. An mRNA Vaccine Protects Mice against Multiple Tick-Transmitted Flavivirus Infections. Cell Rep 25:3382–3392 e3.

58. VanBlargan LA, Errico JM, Kafai NM, Burgomaster KE, Jethva PN, Broeckel RM, Meade-White K, Nelson CA, Himansu S, Wang D, Handley SA, Gross ML, Best SM, Pierson TC, Fremont DH, Diamond MS. 2021. Broadly neutralizing monoclonal antibodies protect against multiple tick-borne flaviviruses. J Exp Med 218.

59. Hinten SR, Beckett GA, Gensheimer KF, Pritchard E, Courtney TM, Sears SD, Woytowicz JM, Preston DG, Smith RP, Jr., Rand PW, Lacombe EH, Holman MS, Lubelczyk CB, Kelso PT, Beelen AP, Stobierski MG, Sotir MJ, Wong S, Ebel G, Kosoy O, Piesman J, Campbell GL, Marfin AA. 2008. Increased recognition of Powassan encephalitis in the United States, 1999-2005. Vector Borne Zoonotic Dis 8:733–40.

60. Dalrymple NA, Mackow ER. 2012. Endothelial cells elicit immune-enhancing responses to dengue virus infection. J Virol 86:6408–15.

61. Sava P, Ramanathan A, Dobronyi A, Peng X, Sun H, Ledesma-Mendoza A, Herzog EL, Gonzalez AL. 2017. Human pericytes adopt myofibroblast properties in the microenvironment of the IPF lung. JCI Insight 2.

62. Baron V, Adamson ED, Calogero A, Ragona G, Mercola D. 2006. The transcription factor Egr1 is a direct regulator of multiple tumor suppressors including TGFbeta1, PTEN, p53, and fibronectin. Cancer Gene Ther 13:115–24.

63. Edagawa M, Kawauchi J, Hirata M, Goshima H, Inoue M, Okamoto T, Murakami A, Maehara Y, Kitajima S. 2014. Role of activating transcription factor 3 (ATF3) in endoplasmic reticulum (ER) stress-induced sensitization of p53-deficient human colon cancer cells to tumor necrosis factor (TNF)-related apoptosis-inducing ligand (TRAIL)-mediated apoptosis through up-regulation of death receptor 5 (DR5) by zerumbone and celecoxib. J Biol Chem 289:21544–61.

64. Kumar SS, Tomita Y, Wrin J, Bruhn M, Swalling A, Mohammed M, Price TJ, Hardingham JE. 2016. High early growth response 1 (EGR1) expression correlates with resistance to anti-EGFR treatment in vitro and with poorer outcome in metastatic colorectal cancer patients treated with cetuximab. Clin Transl Oncol doi:10.1007/s12094-016-1596-8.

65. Tyner JW, Uchida O, Kajiwara N, Kim EY, Patel AC, O’Sullivan MP, Walter MJ, Schwendener RA, Cook DN, Danoff TM, Holtzman MJ. 2005. CCL5-CCR5 interaction provides antiapoptotic signals for macrophage survival during viral infection. Nat Med 11:1180–7.

66. Lepej SZ, Misic-Majerus L, Jeren T, Rode OD, Remenar A, Sporec V, Vince A. 2007. Chemokines CXCL10 and CXCL11 in the cerebrospinal fluid of patients with tick-borne encephalitis. Acta Neurol Scand 115:109–14.

67. Zajkowska J, Moniuszko-Malinowska A, Pancewicz SA, Muszynska-Mazur A, Kondrusik M, Grygorczuk S, Swierzbinska-Pijanowska R, Dunaj J, Czupryna P. 2011. Evaluation of CXCL10, CXCL11, CXCL12 and CXCL13 chemokines in serum and cerebrospinal fluid in patients with tick borne encephalitis (TBE). Adv Med Sci 56:311–7.

68. Pokorna Formanova P, Palus M, Salat J, Honig V, Stefanik M, Svoboda P, Ruzek D. 2019. Changes in cytokine and chemokine profiles in mouse serum and brain, and in human neural cells, upon tick-borne encephalitis virus infection. J Neuroinflammation 16:205.

69. Klein RS, Lin E, Zhang B, Luster AD, Tollett J, Samuel MA, Engle M, Diamond MS. 2005. Neuronal CXCL10 directs CD8+ T-cell recruitment and control of West Nile virus encephalitis. J Virol 79:11457–66.

70. Glass WG, Lim JK, Cholera R, Pletnev AG, Gao JL, Murphy PM. 2005. Chemokine receptor CCR5 promotes leukocyte trafficking to the brain and survival in West Nile virus infection. J Exp Med 202:1087–98.

71. Palus M, Vojtiskova J, Salat J, Kopecky J, Grubhoffer L, Lipoldova M, Demant P, Ruzek D. 2013. Mice with different susceptibility to tick-borne encephalitis virus infection show selective neutralizing antibody response and inflammatory reaction in the central nervous system. J Neuroinflammation 10:77.

72. Liu M, Guo S, Hibbert JM, Jain V, Singh N, Wilson NO, Stiles JK. 2011. CXCL10/IP-10 in infectious diseases pathogenesis and potential therapeutic implications. Cytokine Growth Factor Rev 22:121–30.

73. Pittaluga A. 2017. CCL5-Glutamate Cross-Talk in Astrocyte-Neuron Communication in Multiple Sclerosis. Front Immunol 8:1079.

74. Barnes DA, Huston M, Holmes R, Benveniste EN, Yong VW, Scholz P, Perez HD. 1996. Induction of RANTES expression by astrocytes and astrocytoma cell lines. J Neuroimmunol 71:207–14.

75. Grygorczuk S, Zajkowska J, Swierzbinska R, Pancewicz S, Kondrusik M, Hermanowska-Szpakowicz T. 2006. [Concentration of the beta-chemokine CCL5 (RANTES) in cerebrospinal fluid in patients with tick-borne encephalitis]. Neurol Neurochir Pol 40:106–11.

76. Zhang X, Zheng Z, Liu X, Shu B, Mao P, Bai B, Hu Q, Luo M, Ma X, Cui Z, Wang H. 2016. Tick-borne encephalitis virus induces chemokine RANTES expression via activation of IRF-3 pathway. J Neuroinflammation 13:209.

77. Liu K, Xiao C, Wang F, Xiang X, Ou A, Wei J, Li B, Shao D, Miao D, Zhao F, Long G, Qiu Y, Zhu H, Ma Z. 2018. Chemokine receptor antagonist block inflammation and therapy Japanese encephalitis virus infection in mouse model. Cytokine 110:70–77.

78. Larena M, Regner M, Lobigs M. 2012. The chemokine receptor CCR5, a therapeutic target for HIV/AIDS antagonists, is critical for recovery in a mouse model of Japanese encephalitis. PLoS One 7:e44834.

79. Veerhuis R, Nielsen HM, Tenner AJ. 2011. Complement in the brain. Mol Immunol 48:1592–603.

80. Mehlhop E, Whitby K, Oliphant T, Marri A, Engle M, Diamond MS. 2005. Complement activation is required for induction of a protective antibody response against West Nile virus infection. J Virol 79:7466–77.

81. Ebenbichler CF, Thielens NM, Vornhagen R, Marschang P, Arlaud GJ, Dierich MP. 1991. Human immunodeficiency virus type 1 activates the classical pathway of complement by direct C1 binding through specific sites in the transmembrane glycoprotein gp41. J Exp Med 174:1417–24.

82. Veje M, Studahl M, Bergstrom T. 2019. Intrathecal complement activation by the classical pathway in tick-borne encephalitis. J Neurovirol 25:397–404.

83. Vasek MJ, Garber C, Dorsey D, Durrant DM, Bollman B, Soung A, Yu J, Perez-Torres C, Frouin A, Wilton DK, Funk K, DeMasters BK, Jiang X, Bowen JR, Mennerick S, Robinson JK, Garbow JR, Tyler KL, Suthar MS, Schmidt RE, Stevens B, Klein RS. 2016. A complement-microglial axis drives synapse loss during virus-induced memory impairment. Nature 534:538–43.

84. Saitou N, Nei M. 1987. The neighbor-joining method: a new method for reconstructing phylogenetic trees. Mol Biol Evol 4:406–25.

85. Felsenstein J. 1985. Confidence Limits on Phylogenies: An Approach Using the Bootstrap. Evolution 39:783–791.

86. Kumar S, Stecher G, Li M, Knyaz C, Tamura K. 2018. MEGA X: Molecular Evolutionary Genetics Analysis across Computing Platforms. Mol Biol Evol 35:1547–1549.

87. Sievers F, Wilm A, Dineen D, Gibson TJ, Karplus K, Li W, Lopez R, McWilliam H, Remmert M, Soding J, Thompson JD, Higgins DG. 2011. Fast, scalable generation of high-quality protein multiple sequence alignments using Clustal Omega. Mol Syst Biol 7:539.

